# The chloroplast envelope localization of protoporphyrinogen oxidase 2 prevents full complementation of Arabidopsis *ppo1*

**DOI:** 10.1101/2022.11.22.517550

**Authors:** Boris Hedtke, Sarah Melissa Strätker, Andrea C. Chiappe Pullido, Bernhard Grimm

## Abstract

All land plants encode two isoforms of protoporphyrinogen oxidase (PPO). While PPO1 is predominantly expressed in green tissues and its loss is seedling-lethal in Arabidopsis, the effects of PPO2 deficiency have not been investigated in detail. We identified two *ppo2* T-DNA insertion mutants from publicly available collections, one of which (*ppo2-2*) is a knock-out mutant. While the loss of PPO2 did not result in any obvious phenotype, significant changes in PPO activity were measured in etiolated and root tissues. However, *ppo1ppo2* double mutants are embryo-lethal. To shed light on possible functional differences between the two isoforms, PPO2 was overexpressed in the *ppo1* background. Although the *ppo1* phenotype was partially complemented, even strong overexpression of PPO2 was unable to fully compensate for the loss of PPO1. Analysis of its subcellular localization revealed that PPO2 is found exclusively in chloroplast envelopes, while PPO1 accumulates in thylakoid membranes. A mitochondrial localization of PPO2 in Arabidopsis was ruled out. Since *A. thaliana PPO2* does not encode a cleavable transit peptide, integration of the protein into the chloroplast envelope must make use of a non-canonical import route. However, when a chloroplast transit peptide was fused to the N-terminus of PPO2, the enzyme was detected predominantly in thylakoid membranes, and was able to fully complement *ppo1*. Thus, the two PPO isoforms in Arabidopsis are functionally equivalent, but spatially separated. Their distinctive localizations within plastids thus enable the synthesis of discrete sub-pools of the PPO product protoprophyrin IX, which may serve different cellular needs.

## Introduction

Tetrapyrrole synthesis (TPS) is a fundamental metabolic pathway found in almost all living organisms. In plants, it consists of a series of at least 25 enzymatic reactions, and gives rise to chlorophylls as well as siroheme, heme and phytochomobilin (Tanaka and Tanaka, 2007). While chlorophyll is thought to be the major end-product of the branched pathway in photoautotrophic plant tissue, demand for different tetrapyrroles varies strongly, depending on the cell type and developmental stage considered (Tanaka and Tanaka, 2007). Since most of the late tetrapyrrole intermediates absorb light, and are therefore potentially detrimental, their accumulation must be strictly controlled. Numerous regulatory circuits are involved in TPS. Indeed, control of the initial steps in 5-aminolevulinic acid (ALA) synthesis alone requires several feedback mechanisms. The rate-limiting step in TPS is catalysed by glutamyl-tRNA reductase (GluTR), which is responsible for the first step in the synthesis of ALA. The activity, stability and subplastidal localization of GluTR are regulated by multiple factors that serve to modulate the supply and allocation of ALA for chlorophyll and the other tetrapyrroles (Meskauskiene et al., 2001; Czarnecki et al., 2011; Wang et al., 2018; Richter et al., 2019). In plants, TPS takes place exclusively within plastids. While all reactions, from ALA formation to the synthesis of protoporphyrinogen IX (Protogen), are catalyzed by soluble proteins in the plastidal stroma, all of the following enzymes, starting with protoporphyrinogen oxidase (PPO), are associated with organellar membranes (Joyard et al., 2009). By extracting six electrons from Protogen, PPO forms protoporphyrin IX (Proto), the last common precursor of chlorophyll and hemesynthesis. The genes that encode PPO in plants belong to the HemY family of oxygenic, FAD-containing enzymes (Kobayashi et al., 2014). HemY-type PPOs are targeted by several classes of inhibitors that are agronomically important as herbicides. By blocking substrate binding, these inhibitors cause rapid accumulation of Protogen. The latter is non-specifically oxidized to Proto, which generates singlet oxygen in the presence of light, leading mainly to lipid peroxidation and, ultimately, to cell death (Jacobs et al., 1991) (Lee and Duke, 1994).

All embryophytic plants possess two HemY-type PPO enzymes, and phylogenetic analyses point to a gene duplication event early in the evolution of land plants (Kobayashi et al., 2014). Consequently, in angiosperms, the isoforms PPO1 and PPO2 share only 25% amino-acid-sequence identity (Lermontova et al., 1997). However, based on crystal structures available for *Nicotiana tabacum* PPO2 (Koch et al., 2004), *Bacillus subtilis* (Qin et al., 2010) and *Homo sapiens* (Qin et al., 2011), the overall folding pattern is largely conserved, even over long phylogenetic distances within the HemY family.

Plant PPO1 is translocated into plastids via a cleavable transit peptide at the N-terminus of the pre-protein (Lermontova et al., 1997). Knock-down of *PPO1* results in photodynamic lesions in tobacco antisense lines (Lermontova and Grimm, 2006), while knock-out mutants in Arabidopsis are seedling-lethal (Zhang et al., 2014). The localization of PPO2 is less clear. Initial analyses in tobacco suggested that PPO2 accumulates in mitochondria (Lermontova et al., 1997). Indeed, dual targeting of PPO2 to mitochondria and chloroplasts has been described for spinach (Watanabe et al., 2001) and correlates with a specific N-terminal extension found in members of the *Amaranthaceae* family. This extended N-terminal region was suggested to confer dual targeting by alternative use of two in-frame initiation codons (Watanabe et al., 2001). However, in Arabidopsis, proteome studies have identified PPO2 as a constituent of the plastid envelope membrane (Joyard et al., 2009). While PPO2-deficient plants have not been investigated so far, recent reports on herbicide resistance specifically conferred by mutations in PPO2 have drawn renewed attention to this specific isoform (Porri et al., 2022).

In the present study, we characterized Arabidopsis *ppo2* mutants and analyzed the localization of the PPO2 protein. Based on the complementation of seedling-lethal *ppo1* plants by *PPO2*, we demonstrated that the two PPO isoforms are spatially separated within plastids and assessed the degree of functional equivalence between them.

## Materials and Methods

### Plant material and growth conditions

The Arabidopsis genotypes used included the wild type (Col-0), *ppo1* (GK_539C07), *ppo2-1* (SALK_141571) and *ppo2-2* (SAIL_841_G04). Plants were grown on soil at 23°C and 100 μmol photons m^-2^ s^-1^ under short-day conditions (10 h light, 14 h dark). Seeds used for etiolated growth were stratified for 3 days at 4°C, followed by light induction for 5 h at 23°C. Etiolated seedlings were grown for 4 days at room temperature (RT) either on plates containing Murashige and Skoog medium (4.4 g L^-1^ Murashige and Skoog, 0.05% [w/v] MES, 0.8% [w/v] agar, pH 5.7) or on soil-filled pots covered with gauze.

### Genotyping PCRs

*ppo2-1* (SALK_141571) and *ppo2-2* mutants were tested for T-DNA insertions in *PPO2* using the primer pairs PPO2R2/pROK_LB4 and PPO2R2/Garlic_LB, respectively. The presence of wild-type *PPO2* alleles was analysed in both lines using PPO2F2/PPO2R2. Primer walking used to localize the insertion event in *ppo2-2* employed primers I–IV, in combination with PPO2F3. Analyses of the *ppo1* mutant (GK_539C07) were performed with the primers PPO1 down fw/Gabi LB (T-DNA) and AtPPO1_genot.WtFw/AtPPO1_genot.WtRv (wt allele). Primer sequences used are listed in Supplementary Table 1.

### Cloning

The primer combinations PPO1_Promoter_FW_SacI/PPO1_5’UTR_RV_fusion and PPO1_3’UTR_FW_fusion/PPO1_3’UTR_RV_PmlI were used to amplify the *PPO1* promoter and 3’UTR, respectively, from wild-type *A. thaliana* DNA. The two fragments were fused using the outer primers, and cloned into pJET (Thermo Scientific). The insert was then transferred into the binary vector pCAMBIA 3301 (Cambia, Canberra, Australia) using SacI and PmlI, thus giving rise to pCAM_PPO1, in which the *PPO1* promoter and 3’UTR are separated by AscI and SbfI restriction sites. The wild-type Arabidopsis *PPO2* was amplified as a 3.8-kb genomic fragment using PPO2_FW_AscI/PPO2_RV_SbfI, and ligated into AscI/SbfI-cleaved pCAM_PPO1, resulting in pPPO1:PPO2. To generate p35S:PPO2, *PPO2* was amplified using PPO2_FW_SacI /PPO2_RV_SacI, and ligated into pGL1 (Apitz et al., 2014) after restriction with SacI.

DNA fragments encoding the 37 N-terminal amino acids of PPO1 and PPO2 (lacking the initial methionine) were amplified using the primer pairs PPO1_FW_SacI/P1P2_Fusion_RV and P1P2_Fusion_FW/PPO2_RV_SacI, respectively. Subsequent fusion of the fragments was achieved by annealing and amplification using the outer SacI primers. The product was transferred into pGL1 via SacI cloning and yielded p35S:TP^PPO1^PPO2. Alternatively, the fusion was re-amplified using PPO1_FW_AscI/PPO2_RV_SbfI and inserted into pCAM_PPO1 cleaved with AscI and SbfI, resulting in pPPO1:TP^PPO1^PPO2.

Fusion of PPO2 to the transit peptide of RBCS was achieved by amplification with RbcS_FW_PmlI/RbcsP2_Fusion_RV and RbcsP2_Fusion_FW/ PPO2_RV_SacI. A fusion PCR utilizing outer PmlI/SacI primers and subsequent cloning into SmaI/SacI-cut pGL1 resulted in p35S:TP^RBCS^PPO2.

### qPCR

RNA was extracted from frozen plant material homogenized in a MM 400 ball mill (Retsch, Germany) using the citric-acid protocol (Onate-Sanchez and Vicente-Carbajosa, 2008). Total RNA (1 μg) was transcribed following DNase I treatment using Moloney Murine Leukemia Virus reverse transcriptase and an oligo dT(18) primer according to the manufacturer’s protocol (ThermoFisher Scientific). qPCR analysis was carried out in a CFX96-C1000 96-well plate thermocycler (Bio-Rad, CA) using ChamQ Universal SYBR qPCR master mix (Vazyme). Calculation of gene expression levels was performed with the Bio-Rad CFX-Manager Software 1.6 using the 2^-ΔΔC(t)^ method. Transcript accumulation was normalized using *SAND* (AT2G28390, (Czechowski et al., 2004)).

### Protein extraction and western-blot procedures

Total leaf protein was extracted from homogenized leaf material using 10 μl of 2x Laemmli buffer (Sambrook, 2001) per mg fresh weight. Samples were heated for 5 min at 95°C, centrifuged for 5 min (16,000 g, RT). Samples representing either identical fresh weight (1 mg) or adjusted to Chl/protein content were fractionated by electrophoresis on SDS-polyacrylamide gels (12%) and blotted onto nitrocellulose membranes (Amersham Protran, GE Healthcare, UK). Membranes were stained with Ponceau S and probed with protein-specific antibodies according to (Sambrook, 2001). Polyclonal antisera against His-tagged recombinant Arabidopsis proteins GluTR1, GBP, FLU, PPO2, FC2, CHLI, CHLM, PORB, and DNAJD12 as well as tobacco PPO1 were raised in the authors lab and affinity-purified using the antigen when necessary. Sera specific for LHCB1.6 and TIC110 were purchased from Agrisera (Sweden).

### HPLC analyses

Tetrapyrroles were extracted from homogenized leaves using 300 μl of acetone:0.2 M NH_4_OH (9:1), incubated for 30 min at −20°C and centrifuged for 30 min (16,000x*g*, 4°C). The supernatant was analyzed by HPLC for (Mg) porphyrins and Chls. Non-covalently bound (ncb) heme was extracted from the remaining pellet by resuspension in 200 μl AHD (acetone:HCl:DMSO, 10:0.5:2) and incubated for 15 min at RT. After centrifugation for 15 min (16,000x*g*, RT), the amount of ncb heme in the supernatant was quantified by HPLC. HPLC analyses were performed on Agilent LC systems following the methods described in Richter et al. (2019), using authentic standards for peak quantification.

### Purification of organelles

Chloroplasts were purified from 6-week-old Arabidopsis plants grown under short-day conditions. Aliquots (20 g) of leaf material were homogenized in 450 mM sorbitol, 20 mM Tricine, 10 mM EDTA, 10 mM NaHCO_3_, and 0.1% BSA (pH 8.4) using a modified Waring blender (Kannangara et al., 1977). The homogenate was filtered through Miracloth and pelleted for 8 min at 500x*g*. Following careful re-suspension with a soft brush in RB (300 mM sorbitol, 20 mM Tricine, 2.5 mM EDTA, 5 mM MgCl_2_, pH 8.4), the suspension was loaded on a step gradient of 40% and 80% Percoll in RB. Centrifugation for 30 min at 6,500x*g* in a swing-out rotor enriched for intact chloroplasts at the Percoll interphase. Chloroplasts were carefully transferred, diluted in RB and pelleted for 6 min at 3,800x*g*. Purified chloroplasts were lysed hypotonically in HLB (25 mM HEPES-KOH, pH 8.0) supplemented with protease inhibitor cocktail (Hycultec, Germany). Following incubation on ice for 1 h, 0.7 volumes of a mixture containing 0.6 M sucrose, 25 mM HEPES-KOH (pH 8.0) and 4 mM MgCl_2_ were added and loaded on a sucrose step gradient consisting of 0.4 M, 1.0 M and 1.2 M sucrose, each in 25 mM HEPES-KOH (pH 8.0). Ultracentrifugation at 200,000x*g* for 1 h in a swing-out rotor enriched for chloroplast envelopes at the interphase between 0.6 M and 1.0 M sucrose, while thylakoids formed a tight pellet. The latter was resuspended in HLB, while envelope membranes were transferred, washed in HLB and collected by centrifugation at 48,000x*g* for 1 h.

Mitochondria from 20 g of Arabidopsis leaves were isolated by intense homogenization in a modified Waring blender in EB (0.3 M sucrose, 25 mM K-pyrophosphate, 2 mM EDTA, 10 mM KH_2_PO_4_, 1% PVP-40, 1% BSA, 5 mM cysteine, pH 7.5). Following filtration through Miracloth, the plant debris was extracted twice by grinding in a mortar for 10 min with additional EB. The filtrates were then combined and centrifuged (5 min, 1,700x*g*). The supernatant was centrifuged again (10 min, 20,000x*g*), and the pellet was resuspended in WB (0.3 M sucrose, 10 mM MOPS, 1 mM EDTA, pH 7.2) and homogenized in a Potter-Elvejhem homogenizer. Dilution in WB was followed by centrifugation at 2,500x*g* for 10 min. The supernatant was then pelleted again (10 min, 20,000x*g*). The crude mitochondrial fraction was resuspended in WB and loaded onto a step gradient consisting of 18%, 25% and 50% Percoll in MGB (0.3 M sucrose, 10 mM MOPS, pH 7.2). Centrifugation at 40,000x*g* for 45 min enriched the mitochondria on top of the 50% Percoll phase. Four final washes in WB at 20,000x*g* for 10 min each were applied to obtain purified mitochondria.

### PPO activity assays

Plant material was either homogenized under liquid N_2_ in a mixer mill MM400 (Retsch, Germany) or obtained from organelle purifications and dissolved in assay buffer (AB) containing 0.5 M Bis-Tris pH 7.5, 4 mM DTT, 2.5 mM EDTA and 0.004% Tween on ice. Aliquots (50 μl) of the extract were mixed with 150 μl of AB containing 4 μM Protogen at room temperature and Proto formation was recorded over a period of 20 min on a Hitachi F-700 fluorescence spectrophotometer (excitation 405 nm, emission 635 nm). Conversion of relative fluorescence values is based on an authentic Proto standard dissolved in AB.

## Results

Numerous T-DNA insertion mutant lines were tested for *PPO2* knockdown or knockout mutations. Two lines (SALK_141571 and SAIL_841_G04) were confirmed to harbour T-DNA left-border (LB) sequences integrated into different sites within the 201-bp intron 13 of AtPPO2 (Figure 1A). Closer inspection of the integration site in the mutant *ppo2-1* (SALK_141571) localized the LB and right border (RB) sequences of the integrated T-DNA at 65 and 55 bp, respectively, from the 3’ end of exon 13. Hence, the insertion resulted in a small deletion in intron 13. In the case of *ppo2-2* (SAIL_841_G04), analysis of the *PPO2*-specific sequences flanking the T-DNA also localized its LB to intron 13, at a point 33 bp downstream of exon 13 (Figure 1B). Homozygous plants were identified for both *ppo2* mutant lines (Figure 1C). Nevertheless, numerous attempts to amplify the sequence flanking the RB of the T-DNA in the *ppo2-2* line failed. Since the integrity of exon 13 in this mutant was crucial for its further use in functional studies of PPO2, additional PCRs were performed on *ppo2-2* genomic DNA using four different primers (named I–IV; Figure 1B) to pin down the precise insertion site of the RB (Figure 1D). PCRs employing primers I and II failed to amplify specific fragments from *ppo2-2* genomic DNA, while control templates (Col-0 and *ppo2-1*) gave rise to the expected products (Figure 1D, left panels). Since amplicons starting at primers III and IV are intact in *ppo2-2* (Figure 1D, right panels), we concluded that the T-DNA insertion in this line had resulted in the deletion of at least 12 bp of exon 13, in addition to the loss of 33 bp of intron 13.

**Figure 1.**
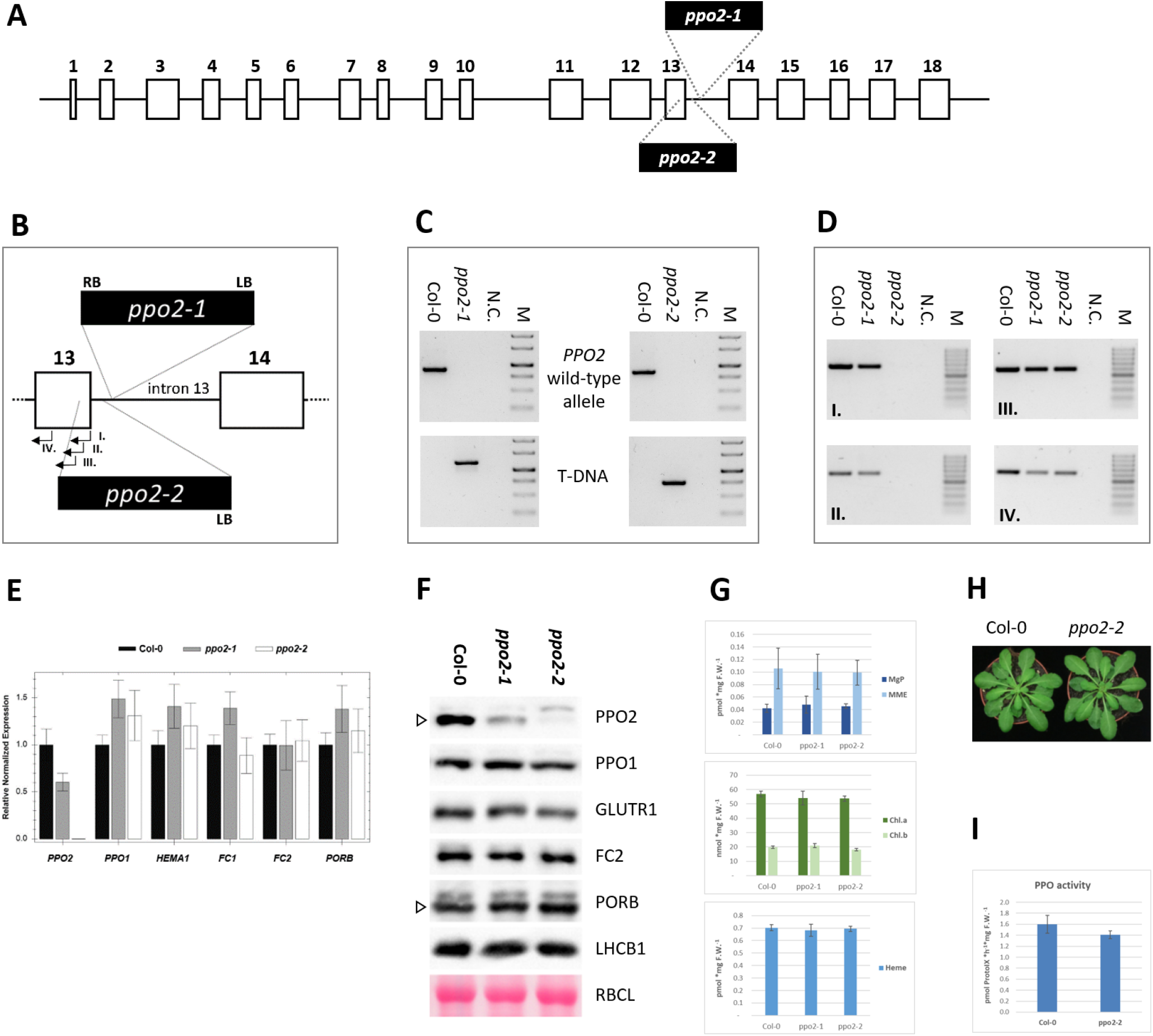
Characterization of Arabidopsis *ppo2* mutants. **A,** Structure of the Arabidopsis *PPO2* gene. Boxes and numbers indicate the exons of the coding region. **B,** Schematicm depiction showing details of the insertions in intron 13 in the mutants *ppo2-1* and *ppo2-2*. Sites of integration of LB and RB sequences are indicated by dashed lines. Exons 13 and 14 are boxed. Primers used for mapping *ppo2-2* (see D.) are indicated by arrows and Roman numerals I-IV. **C,** Genotyping PCRs confirm the homozygosity of T-DNA insertions in *ppo2-1* and *ppo2-2*, respectively. Pairs of gene primers specific for the wild-type *PPO2* allele (upper panel) or a combination of gene- and LB-specific primers that amplify only *PPO2* alleles interrupted by T-DNA integration (lower panel) were used. **D,** PCR test of the integrity of the coding region of exon 13 in *ppo2-2*. Four different primers that bind to sequences in exon 13 (labeled I-IV; see also **B**) were used to map the deletion caused by T-DNA integration in *ppo2-2*. Wild-type (Col-0) and *ppo2-1* strains served as controls. Negative controls (N.C.) were performed without added DNA template. GeneRuler 1 kb (C) or 100 bp (D) DNA ladders (Thermo Scientific) were used as markers (M). **E,** Quantification of *PPO2, PPO1*, GluTR1 (*HEMA1), FC1, FC2* and *PORB* transcripts in *ppo2* mutants relative to wild type. Quantitative real-time PCR data were normalized to SAND transcript accumulation. Seedlings were cultivated for 14 days under short-day (SD) conditions. **F,** Quantification of the PPO2 protein, selected tetrapyrrole pathway enzymes and light-harvesting complex protein LHCB1. As a reference, Ponceau-stained RBCL (RuBisCO large subunit) is depicted. Where antisera show cross-reactions, the specific band is marked by an open triangle. **G,** Quantification of TPS intermediates (MgP, MME) and the end-products chlorophyll (shown separately as Chl.a and Chl.b) and heme in 4-week-old *ppo2* mutants and the wild type. **H,** Phenotype of *ppo2-2* plants after 6 weeks of cultivation under SD conditions. **I,** Comparison of PPO activity in leaf extracts of wild-type and *ppo2-2 m* utant plants. Enzymatic activity is expressed as the amount of Proto formed per unit time and fresh weight of leaf material. The assay used 2-week-old seedlings grown under SD conditions.

PPO2 transcript abundance in both *ppo2* T-DNA insertion lines was evaluated by quantitative real-time PCR (qPCR). While the T-DNA insertion in intron 13 of line *ppo2-1* results in a 40% decrease in *PPO2* transcripts, no *PPO2* mRNA at all is detectable in *ppo2-2* (Figure 1E). Transcript abundances of *PPO1* and selected key TPS genes (*HEMA1* encoding glutamyl tRNA reductase1 (GluTR1), *FC1* and *FC2* for the two ferrochelatase isoforms, *PORB* for protochlorophyllide oxidoreductase B) were not affected in either of the *ppo2* mutants.

Protein levels were characterized by immunoblot analyses (Figure 1F). Antisera raised against AtPPO2 revealed significantly reduced amounts of the PPO2 protein in *ppo2-1*. In protein extracts of the *ppo2-2* mutant, no PPO2-specific signal was detected. Immunoblot analyses of TPS- and photosynthesis-related proteins revealed no significant change in amounts of PPO1, GluTR1, FC2, PORB or the light-harvesting complex protein B1 (LHCB1) in either of the *ppo2* mutants in comparison to wild type (Figure 1F).

Since both genomic and expression analyses characterized *ppo2-2* as a knockout mutant, further characterization was focused on this line. Analyses of the metabolic intermediates Mg protoporphyrin IX (MgP), Mg protoporphyrin monomethylester (MME), as well as their end-products chlorophyll and heme, indicated no significant disturbance of the TPS pathway under standard growth conditions in *ppo2-2* (Figure 1G). Moreover, the visible phenotype of *ppo2-2* plants did not differ from wild type (Figure 1H).

In order to evaluate the contribution of PPO2 to total plant PPO activity under photoautotrophic conditions, the level of PPO enzymatic activity in leaf extracts of *ppo2-2* was compared to that in wild-type samples. In 2-week-old seedlings, total PPO activity in *ppo2-2* was reduced by 12% in comparison to Col-0 (Figure 1I). This relatively minor impact of the loss of PPO2 on total plant PPO activity is in good agreement with relative transcript abundances described for the two PPO isoforms in Arabidopsis (Winter et al., 2007). In the majority of developmental stages, in green photoautotrophic tissue and in most plant organs, *PPO1* transcripts are clearly dominant over their *PPO2* counterparts.

Thus, in light of the markedly unequal contributions of the two isoforms to PPO activity, the divergent phenotypic consequences of the two knockout mutants are plausible. While loss of *PPO1* function is seedling-lethal (Zhang et al., 2014), Figure 2A), the *ppo2-2* knockout mutant described here for the first time displays wild-type-like growth. Interestingly, *ppo1* plantlets also do not differ from wild type when cultivated under etiolating conditions (Figure 2B). This suggests that PPO2 activity makes a significantly larger contribution to TPS during germination and development in the dark.

**Figure 2.**
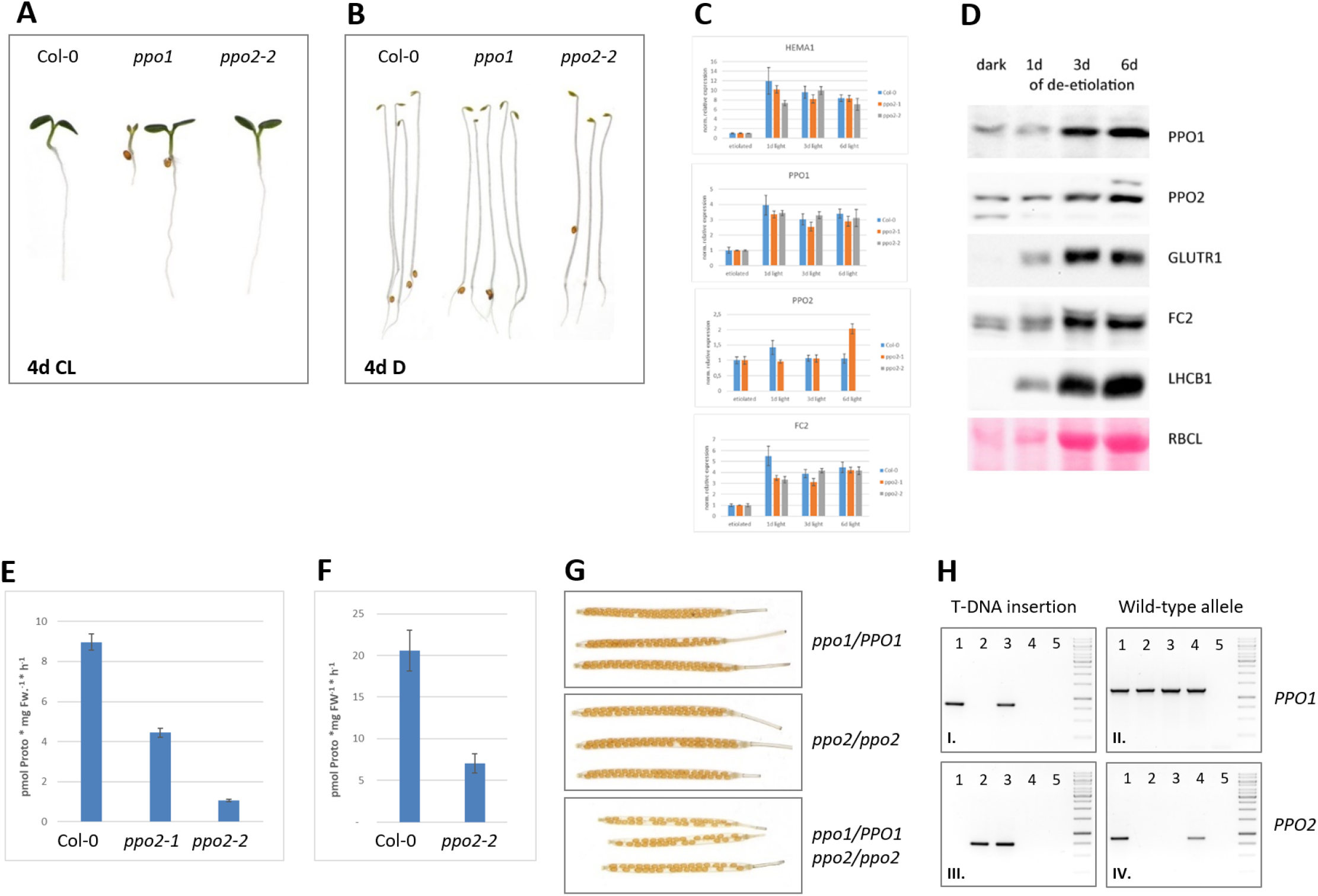
Contribution of the PPO2 isoform in etiolated seedlings, roots and seed development. **A,** Comparison of wild-type and *ppo* mutant phenotypes after 4 days in continuous light. *ppo1* seedlings were derived from a heterozygous parent and segregate. **B,** Wild-type and *ppo* mutant seedlings after 4 days growth in darkness. **C,** Changes in accumulation of *HEMA1, PPO1, PPO2* and *FC2* transcripts after exposure of 4-day-old etiolated seedlings to light for 1, 3 and 6 days. Analyses were performed for wild-type (Col-0), *ppo2-1* and *ppo2-2* seedlings. mRNA accumulation is given relative to etiolated (=1) seedlings, expression was normalized to accumulation of SAND. Standard deviation are based on 3 biological replicates. **D,** Immunoblot analysis of PPO, PPO2, GluTR1, FC2 and LHCB1 levels during de-etiolation. The Ponceau-stained membrane (lower panel) depicts the accumulation of RBCL at the investigated time points. **E,** PPO enzyme activity in etiolated wild type (Col-0) in comparison to *ppo2-1* and *ppo2-2*. Total extracts of seedlings grown for 4 days in darkness were used to quantify Proto formation *in vitro*. **F,** PPO enzyme activity in roots of wild type (Col-0) relative to *ppo2-2*. Seedlings were grown for 15 days in continuous light on MS plates kept in a vertical position. Roots were then cut and used to measure Proto formation *in vitro*. **G,** Crosses of heterozygous *ppo1* mutants (*ppo1/PPO1*) to homozygous *ppo2-2* plants (*ppo2/ppo2*) gave rise to *ppo1/PPO1 ppo2/ppo2* individuals in generation F2. Siliques formed by these plants show an abortion rate equivalent to approximately 25% of the developing seeds (lower panel). For comparison, siliques of the crossed lines are depicted in the upper panels. **H,** Genotyping PCR analyses of plants whose seed formation is shown in G. Plants heterozygous for *ppo1* (1), homozygous for *ppo2-2* (2), together with their *ppo1/PPO1 ppo2/ppo2* F2 siblings (3), are shown next to wild type (4) and non-template controls. Tests for inserted T-DNA in *PPO1* (I.) and *PPO2* (III.) are shown next to PCRs confirming the integrity of wild-type alleles of *PPO1* (II.) and *PPO2* (IV.). A DNA marker (GeneRuler 1 kb, ThermoFisher) is shown on the right.

To verify the more prominent role of PPO2 under these conditions, transcripts encoding both PPO isoforms were quantified in wild-type seedlings after 4 days of growth under etiolating conditions and during subsequent de-etiolation (Figure 2C). Levels of *PPO1* transcripts during de-etiolation reveal light-inducible expression, which resembles that of both *HEMA1* and *FC2* (Figure 2C). In contrast, amounts of *PPO2* mRNA are not significantly influenced by light. This difference in light-responsiveness between the two PPO isoforms was also observed at the protein level (Figure 2E). While levels of PPO1 are clearly enhanced after three days of illumination, PPO2 abundance is significantly less affected.

Due to the divergent responses of the PPO isoforms during de-etiolation, we also assessed the contribution of PPO2 to total PPO activity in dark-grown seedlings. In contrast to the minor decrease observed under photoautotrophic conditions (Figure 1I), *in-vitro* PPO activity is reduced by 88% in etiolated *ppo2-2* plantlets relative to wild type (Figure 2E). In agreement with this marked decrease in PPO activity in *ppo2-2*, knockdown of *PPO2* expression in *ppo2-1* results in a clearly reduced PPO activity in etiolated seedlings (Figure 2E). Since dark-grown seedlings show a stronger contribution of PPO2 activity in non-photoautotrophic tissues, PPO activity was also measured in roots of *ppo2-2* mutants. In comparison to wild-type tissue, *ppo2-2* roots revealed a 66% decrease in PPO activity (Figure 2F).

Although the higher impact of PPO2 on overall PPO activity in etiolated seedlings and roots does not result in a visible phenotype, the results hint at a substantial contribution of *PPO2* during germination and early seedling growth. Moreover, since the *ppo1* mutant develops like the wild type in the dark (Figure 2B), TPS during embryo and seed development is presumably supported predominantly by PPO2 function. To test this assumption, *ppo1ppo2* double mutants were generated. Crosses of heterozygous *ppo1* to homozygous *ppo2-2* enabled the identification of individuals heterozygous for *ppo1* and homozygous for *ppo2-2 (ppo1/PPO1 ppo2/ppo2*) among the F2 offspring. These plants were phenotypically indistinguishable from wild type. However, analysis of seed development in ripening siliques revealed that around 25% of silique positions were empty. This corresponds to the expected number of seeds that were homozygous for the *ppo1ppo2* double mutation (Figure 2G). No significant abortion of seeds was observed in siliques of the heterozygous *ppo1* mutants used in this study (GABI_539C07) or in homozygous *ppo2-2* mutants, i.e. the parental lines of the cross (Figure 2G). PCR analyses confirmed the respective genotypes (Figure 2H).

### Complementation of *ppo1* by overexpression of PPO2

Since PPO1 is the dominantly expressed isoform in light-exposed leaves, seedling lethality of *ppo1* is likely to be caused by a reduction in overall PPO activity which results in the accumulation of phototoxic Proto. To assess the functional equivalence of the two PPO isoforms, as well as the consequences of an increased contribution of PPO2 in green tissue, *PPO2* was expressed in the *ppo1* mutant background under the control of the *PPO1* promoter (*pPPO1*). Transformation of heterozygous *ppo1* mutants enabled the identification of BASTA-resistant T1 seedlings exhibiting enhanced *PPO2* expression in a heterozygous *ppo1* mutant background. Quantification of *PPO2* mRNA levels in three representative transgenic lines revealed a 5- to 10-fold increase in *PPO2* transcript levels compared to wild type, while amounts of the *PPO1* transcript were slightly reduced due to *ppo1* heterozygosity (Figure 3A, lower left and middle). Since the *pPPO1:PPO2* DNA construct was designed to incorporate the untranslated regions (UTRs) of *PPO1* (upper scheme in Figure 3A), qPCR quantification using primers specific for the 3’UTR of *PPO1* included *PPO1* mRNA as well as transgenic *PPO2* transcripts, thus enabling quantitative comparisons of transgene expression with endogenous *PPO1* mRNA levels. The observed 5- to 10-fold increase in *PPO2* mRNA levels in the three depicted lines corresponded to an estimated 2-fold increase in amounts of the *PPO1* 3’UTR, when the slightly lower endogenous PPO1 expression is considered in heterozygous lines of the T1 offspring (Figure 3A, right panel). Hence, as expected for expression under control of *pPPO1*,transgenic *PPO2* is expressed in similar amounts to wild-type *PPO1*.

**Figure 3.**
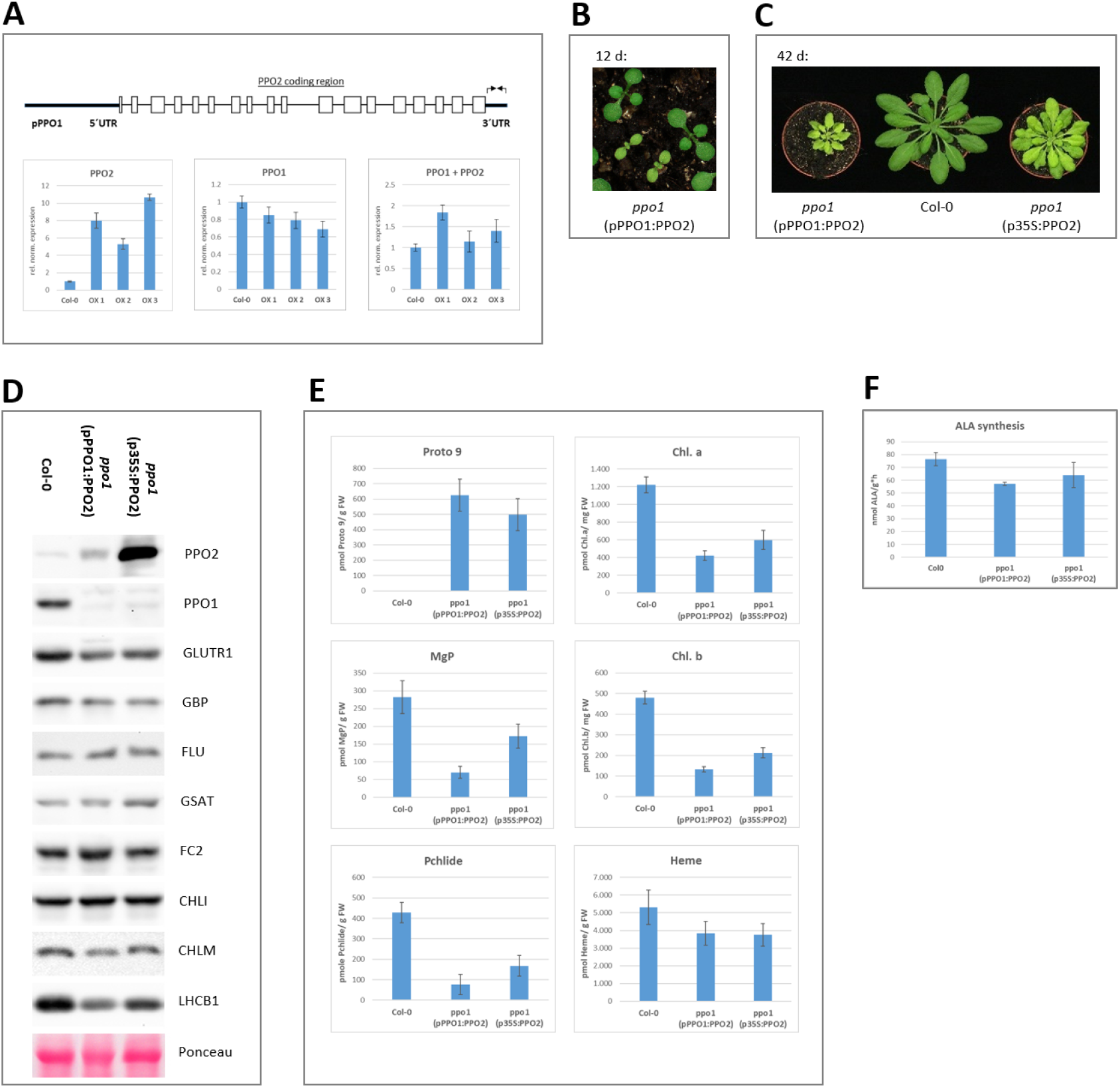
Overexpression of *PPO2* complements the *ppo1* phenotype only partially. **A,** Upper panel: Schematic depiction of the construct pPPO1:PPO2. Besides the *PPO1* promoter (pPPO1), the 5’UTR and the 3’UTR of *PPO1* (bold lines) flank the genomic sequence of *PPO2*. Exons are boxed. Primers specific for the 3’UTR of *PPO1* are indicated as arrows. Lower panel: qPCR analysis of transgenic plants expressing pPPO1:PPO2. Three lines (OX1-3) were analyzed in comparison to wild type (Col-0) for expression of *PPO2, PPO1* as well as the 3’UTR of *PPO1* (“PPO1+PPO2”), which quantifies *PPO1* as well as the transgenic *PPO2*. Transcript abundance was normalized using expression of SAND. **B,** Phenotypes of representative 12-day-old T2 seedlings of *ppo1* expressing with pPPO1:PPO2. **C**, Phenotype of *ppo1* mutants complemented by expression of PPO2 under control of pPPO1 (left) or p35S (right) in comparison to wild type. All depicted individuals were grown for 6 weeks under short-day conditions. **D,** Immunoblot analysis of selected TPS enzymes (PPO1, PPO2, GluTR1, GBP, FLU, GSAT, FC2, CHLI, CHLM) and light-harvesting proteins (LHCB1). The Ponceau-stained membrane depicts the accumulation of RBCL in the analyzed samples. **E,** Accumulation of tetrapyrrole intermediates (Proto, MgP, Pchlide) and end-products (Chl.a, Chl.b, heme). **F,** ALA-synthesizing capacity was determined in the presence of levulinate to inhibit ALAD activity.

Inspection of the T2 offspring of heterozygous *ppo1(pPPO1:PPO2*) individuals revealed seedlings with a deviating phenotype: about one quarter of T2 seedlings displayed pale green cotyledons and true leaves in combination with retarded growth (Figure 3B). Aberrant pigmentation and growth were observed during the entire life cycle of these seedlings, as exemplified for 6-week-old mutant seedlings in comparison to wild-type plants (Figure 3C, left and middle). Following transfer of adult pale-green mutant plants to continuous light, they flowered and set seeds. T3 siblings of such individuals displayed a uniform phenotype identical to that of their parental T2 plants.

Since expression of *PPO2* under the control of the *PPO1* promoter only partially complemented the *ppo1* phenotype, overexpression of *PPO2* in *ppo1* was further increased by employing the cauliflower mosaic virus (CaMV) 35S promoter (p35S). The introduction of *p35S:PPO2* into the *ppo1* mutant background gave rise to transgenic individuals that complemented the *PPO1* knockout more effectively than lines harbouring pPPO1:PPO2, but were still strongly perturbed in pigmentation and development in comparison to wild type (Figure 3C, right). Transgenic lines expressing *p35S:PPO2* accumulated up to 100 times more *PPO2* transcript in comparison to wild type and similarly excessive amounts of PPO2 protein. Thus, the amounts of PPO2 in these lines exceeded the average PPO1 accumulation in wild type by more than an order of magnitude.

Homozygous *ppo1* mutants carrying the two different *PPO2* expression constructs were subjected to analyses of protein expression and the content of tetrapyrrole metabolites. Immunoanalysis confirmed the pale green mutants as homozygous *ppo1* individuals overexpressing PPO2 to different extents (Figure 3D). Immunoblot analyses of further TPS and LHC proteins revealed significantly reduced contents of GluTR1 and LHCB1 in *PPO2*-complemented *ppo1* plants (Figure 3D). In contrast, amounts of the TPS enzymes GluTR-Binding Protein (GBP), FLUORESCENT (FLU), FC2, Magnesium chelatase subunit I (CHLI), PORB and Magnesium-protoporphyrin IX Methyltransferase (CHLM) were not altered in the partially complemented *ppo1* mutants (Figure 3D). The partial rescue of *ppo1* was underlined by the analysis of TPS intermediates and end-products in the chlorotic mutants. The amount of Proto, which actually represents the sum of Protogen and Proto (owing to auto-oxidation of Protogen during sample extraction and processing), is strongly increased in both complemented lines, while downstream intermediates of the chlorophyll branch, such as MgP and protochlorophyllide (Pchlide) are strongly reduced (Figure 3E). Chlorophyll contents are reduced by 50 to 70% in comparison to wild type, while the concentration of non-covalently bound heme corresponds to 73% of wild-type content, indicating that heme synthesis is less severely affected. The observed decrease in GluTR1 accumulation prompted us to compare ALA synthesis rates. A reduction of ALA synthesis by 18 to 25% was observed in the two complemented mutant lines relative to wild type (Figure 3F).

### Subplastidal localization of PPOs

As even excessive accumulation of PPO2 fails to fully complement the loss of PPO1 activity, the intracellular localization of the two PPO isoforms in Arabidopsis was re-examined. As PPO2 overexpression partially complements the seedling-lethality of *ppo1*, exclusively mitochondrial targeting of PPO2 can be ruled out for Arabidopsis. To provide experimental evidence for the subcellular localization of PPO2 in *A. thaliana*, purified mitochondria and chloroplasts were analyzed.

Firstly, mitochondria purified from wild-type *A. thaliana* plants were examined and selected proteins were analysed on immunoblots (Figure 4A). A strong increase in the signal intensity for the mitochondrial marker protein Voltage-Dependent Anion-selective Channel (VDAC), which belongs to a family of mitochondrial outer-membrane porins, confirmed that the mitochondrial fraction had been successfully enriched (Figure 4A). Nevertheless, PPO2 is detectable only in the total protein fraction. As expected, TIC110, a component of the Translocon at the Inner Chloroplast membrane (TIC) complex, which served as a chloroplast marker protein, is strongly depleted in the mitochondrial fraction (Figure 4A). To underline this finding, the experiment was repeated using transgenic Arabidopsis plants containing p35S:PPO2 to ensure that the protein is efficiently expressed (Figure 4B). The organellar markers VDAC and TIC110 used as controls were distributed as in the wild type. PPO2 was strongly overexpressed, as indicated by the presence of a weak PPO2-specific band in the mitochondrial fraction. However, comparison to the distribution obtained for TIC110 reveals that the mitochondrial fraction contains a contamination of chloroplasts. Residual impurities of plant mitochondrial preparations by plastidal or peroxisomal membranes are unavoidable and have been repeatedly described (Tran and Van Aken, 2022).

**Figure 4.**
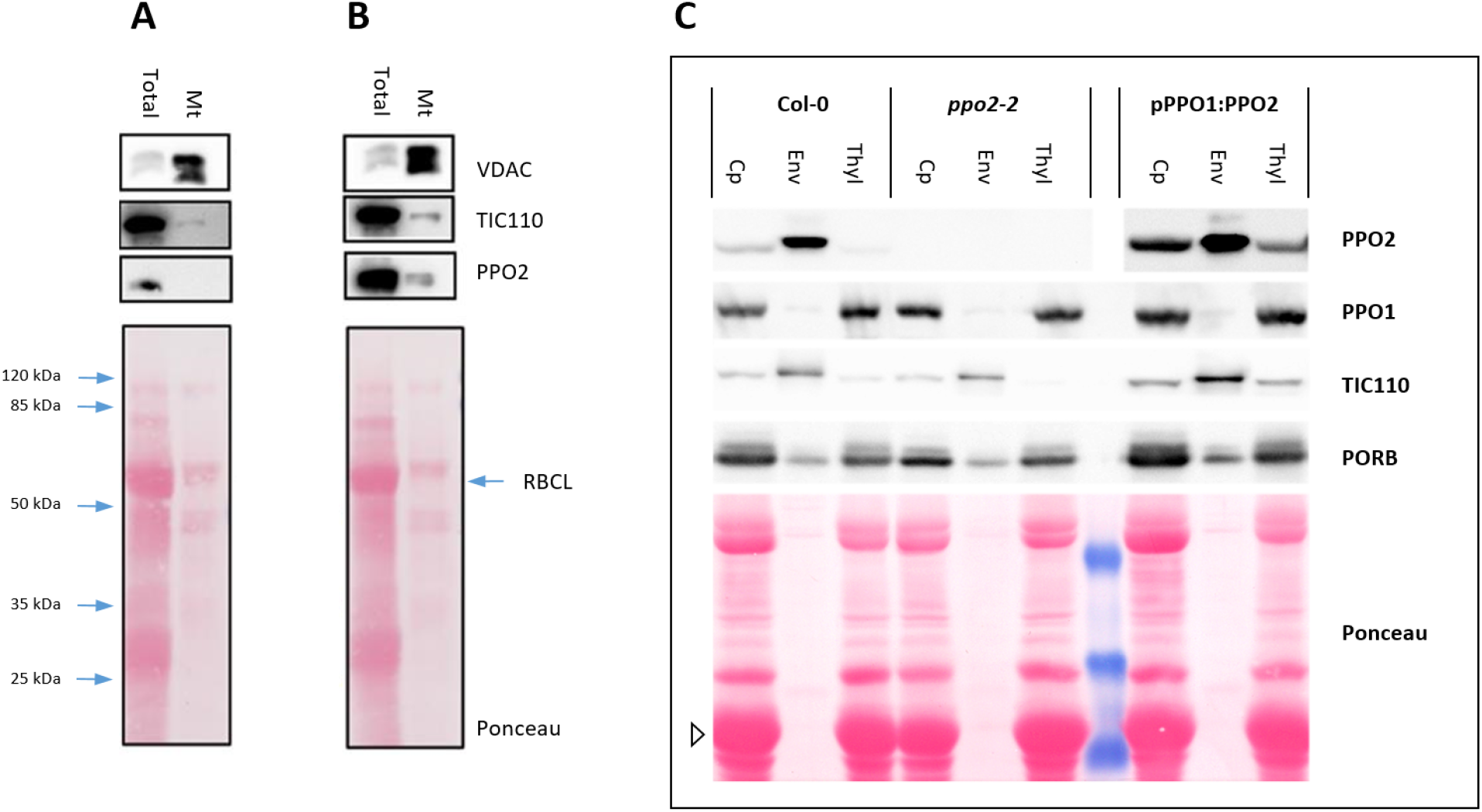
Localization of the two PPO isoforms of *A. thaliana* in different chloroplast membranes. **A-B,** Mitochondria were enriched from wild-type Col-0 (**A**) and PPO2-overexpressing plants (**B**). Total leaf proteins (Total) are loaded next to purified mitochondrial fractions (Mt). Immunoblot analyses were performed using antibodies directed against the mitochondrial voltage-driven anion channel (VDAC), the chloroplast envelope protein TIC110 and PPO2. **C,** Percoll-purified chloroplasts (Cp) from Arabidopsis leaves were lysed hypotonically and used to enrich for plastidal envelopes (Env) and thylakoid (Thyl) membrane fractions by sucrose-density gradient ultracentrifugation. The analyses were performed on the wild type (Col-0), PPO2 knockout mutants (*ppo2-2*) and plants expressing *PPO2* under the control of pPPO1 (pPPO1:PPO2). Immunoblots were incubated with antibodies specific for PPO2, PPO1, TIC110 and PORB. The Ponceau stained membrane/gel includes prestained protein marker bands of 50, 35 and 25 kDa (blue). LHCB1 is indicated by an open triangle.

Having excluded a mitochondrial localization for PPO2, the distribution of the two Arabidopsis PPO isoforms within plastids was addressed (Figure 4C). Leaf chloroplasts were purified on Percoll gradients and, following hypotonic lysis, further separated into envelope and thylakoid fractions using sucrose-gradient ultracentrifugation. Further analysis revealed that PPO1 and PPO2 are differentially distributed in wild-type chloroplasts (Figure 4C, left). While PPO1 is exclusively found in the thylakoid fraction, PPO2 accumulates specifically in envelope membranes. To verify the success of fractionation, proteins of known localization were detected in parallel (Figure 4C). While TIC110 is a chloroplast envelope-specific protein, PORB is known to be enriched in thylakoid membranes. LHCB1 as an additional thylakoid marker is highlighted in the Ponceau staining pattern in Figure 4C.

To validate the specificity of the observed PPO2 localization, the *ppo2-2* knockout mutant as well as an overexpression line was included in the analysis. When chloroplasts from *ppo2-2* plants were fractionated, no PPO2-specific immune signals were detected (Figure 4C, middle). In contrast, plants overexpressing *PPO2* under the control of pPPO1 showed a wild-type-like distribution of PPO2, i.e., enrichment of the transgenic protein in the envelope fraction (Figure 4C, right).

Comparison of PPO activities in purified chloroplast and envelope fractions of wild-type Arabidopsis plants revealed a strong increase in the concentration of Proto present in the envelope-enriched fraction (Col-0 in Table 1). This enhanced activity in purified envelopes reflects the minor contribution of envelopes to total chloroplast proteins: Since only about 1-2% of all chloroplast proteins are localized in the envelope (Douce and Joyard, 1990), a specific enrichment of this fraction results in over-representation of envelope components when comparisons are normalized to protein amounts. As expected, the high level of envelope-localized PPO activity found in wild-type plants was not detectable in *ppo2-2* mutants (Table 1). PPO activity in whole isolated chloroplasts was also lower in *ppo2-2*. In addition, *in-vitro* formation of Proto was investigated using plastidal fractions of *pPPO1:PPO2*-overexpressing plants. Here, a 12-fold increase in PPO activity was observed in purified chloroplasts, most of which was attributable to the envelope fraction (Table 1).

**Table 1.** PPO enzyme activity in suborganellar fractions. Purified chloroplasts (Cp) and purified plastid envelopes purified from wild-type (Col-0), *ppo2-2* and transgenic lines overexpressing PPO2 under the control of pPPO1, and used for *in-vitro* PPO activity assays. The resulting activities are given in pmol of ProtoIX formed per μg protein per min. The standard deviations of three different measurements are indicated in the column on the right.

| Plant line | fraction | PPO activity<br>[pmol Proto*min<sup>-1</sup><br>*µg<sup>-1</sup> protein] | Std. deviation<br>[pmol Proto*min<sup>-1</sup><br>*µg<sup>-1</sup> protein] |
| --- | --- | --- | --- |
| Col-0 | purified Cp | 0.67 | 0.17 |
|  | envelope | 29.61 | 1.19 |
| *ppo2-2* | purified Cp | 0.33 | 0.03 |
|  | envelope | 0.30 | 0.28 |
| pPPO1:PPO2<br>overexpresser | purified Cp | 8.33 | 0.69 |
|  | envelope | 273.59 | 16.94 |

### Arabidopsis PPO2 is strictly confined to the plastid envelope

The evidence for the specific localization of *A. thaliana* PPO2 in chloroplast envelope membranes presented above raised the question of the mechanism of its import. An N-terminally extended amino-acid sequence was previously reported for spinach PPO2 (Watanabe et al., 2001). Among dicotyledonous plants, this amino-terminal extension is shared only by members of the *Amaranthaceae* family (shown for spinach and *Amaranthus palmeri* in Figure 5A). Interestingly, it also seems to be a common feature of PPO2 isoforms in monocotyledonous plants (depicted for maize and rice in Figure 5A). Like the vast majority of dicotyledonous PPO2 sequences, Arabidopsis PPO2 has a different and significantly shorter amino-terminus, which resembles that of the tobacco PPO2 sequence (Figure 5A). At5g14220, the gene encoding Arabidopsis PPO2, gives rise to a transcript containing a 5’ untranslated region (5’ UTR) of 58 nucleotides (At5g14220.1, Araport 11). Since the annotated mRNA does not include an in-frame stop codon upstream of the translation initiation site, the presence of an even longer 5’UTR was excluded by RT-PCR analyses.

**Figure 5.**
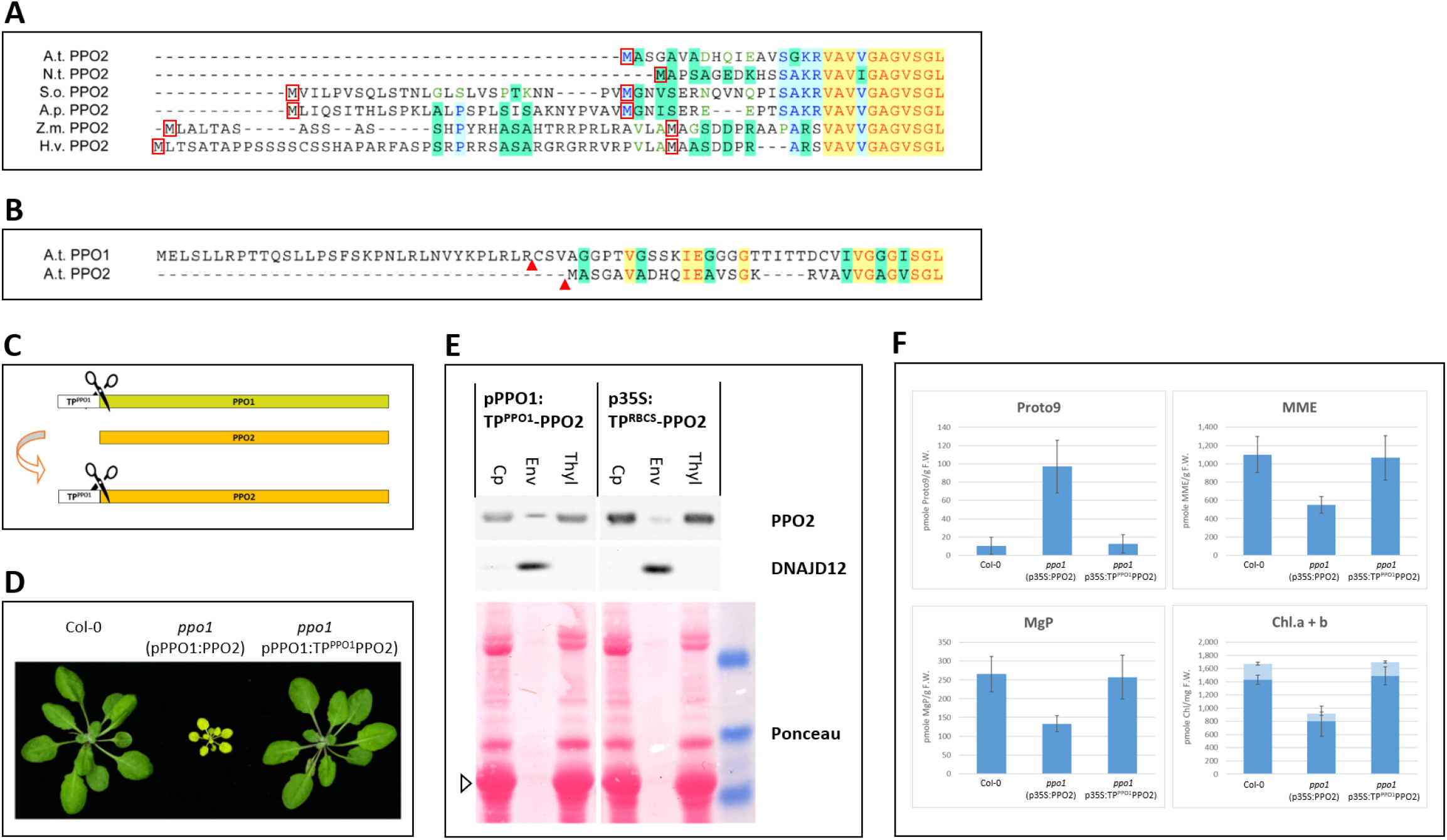
Targeting of PPO2 to thylakoid membranes enables full complementation of *ppo1*. **A**, Alignment of the N-terminal sequences of PPO2 proteins from representative angiosperms. In the majority of dicotyledonous plants, the N-terminus of PPO2 lacks the predicted cleavable transit peptide, as depicted for *A. thaliana* (A.t.) and *N. tabacum* (N.t.). In plants of the *Amaranthaceae* family, as shown here for *Spinacia oleracea* (S.o.) and *Amaranthus palmeri* (A.p.), the N-terminal sequence is extended, as it is in monocotyledonous angiosperms such as *Zea mays* (Z.m.) and *Hordeum vulgare* (H.v.). Red boxes highlight methionine residues. Identical residues are shown in red against a yellow background, blocks of similar amino acids are shaded in turquoise. Conservative exchanges are shown in blue/light blue, weakly similar residues in green. **B,** Alignment of the amino-termini of PPO1 and PPO2 in *A. thaliana*. N-termini of mature PPO proteins identified by mass spectrometry (NTerdb) are indicated by red arrows.**C,** Scheme illustrating the fusion of 37 N-terminal amino acids of A.t. PPO1 to AtPPO2, which gives rise to TP^PPO1^PPO2. **D,** Phenotypic comparison of wild-type and homozygous *ppo1* mutants complemented with pPPO1:PPO2 and pPPO1:TP^PPO1^PPO2, respectively. Plants were cultivated for 30 days under SD conditions (8 h light/16 h dark) and 100 μE light intensity. **E,** Immunoblot analyses of purified chloroplasts (Cp) and fractions enriched for envelope (Env) and thylakoid (Thy) membranes of Arabidopsis plants expressing pPPO1:TP^PPO1^PPO2 and p35S:TP^RBCS^PPO2, respectively, in the Col-0 background. Antisera specific for PPO2 and DNADJ12 were used. The lower panel depicts marker bands of 50, 35 and 25 kDa, prestained with Ponceau Red (right side); LHCB1 is highlighted by an open triangle. **F,** HPLC analyses of *ppo1* complemented with either envelope-(pPPO1:PPO2) or thylakoid-targeted (TP^PPO1^PPO2) PPO2 constructs. The tetrapyrrole intermediates ProtoIX, MgP and MME, as well as Chl a and Chl b, were quantified in 3.5-week-old wild-type and complemented mutant seedlings. Standard deviations for three biological replicates are indicated.

Interestingly, according to NTerdb (https://n-terdb.i2bc.paris-saclay.fr), a compilation of the results of targeted mass-spectrometric analyses of the N-termini of mature *A. thaliana* proteins (Bienvenut et al., 2012), mature PPO2 begins with an acetylated alanine residue that is derived from the second amino acid of the annotated polypeptide (Figure 5B). Since methionine excision and N-terminal acetylations are frequently observed co-translational events (Giglione et al., 2015), the localization of PPO2 to chloroplast envelopes in Arabidopsis does not result from a canonical import event mediated by a cleavable transit peptide (Li and Teng, 2013).

In contrast to the conflicting cellular localizations described for plant PPO2 isoforms in previous studies, data on organellar targeting of PPO1 are consistent. Plant PPO1 sequences harbour a characteristic chloroplast transit peptide and cleavage of 34 N-terminal amino acids has been confirmed in *A. thaliana* by mass spectrometric analysis (https://n-terdb.i2bc.paris-saclay.fr). The mature PPO1 protein is associated with thylakoid membranes (see Figure 4C).

To investigate the degree of functional equivalence between the two Arabidopsis PPO isoforms, the cleavable transit peptide of PPO1 (TP^PPO1^) was fused to the N-terminus of PPO2. The resulting gene construct encoding TP^PPO1^PPO2 included the 37 amino-terminal residues of the PPO1 preprotein (Figure 5C). Since the N-terminal methionine is absent in mature *A. thaliana* PPO2, it was omitted in the TP^PPO1^PPO2 fusion in order to rule out an alternative initiation of translation. The newly generated *TP^PPO1^PPO2* fusion gene was placed under control of *pPPO1* as well as *p35S* and transformed into the (heterozygous) *ppo1* mutant background.

Analyses of BASTA-resistant seedlings identified transformants homozygous for *ppo1* in the T1 generation. The phenotype of *ppo1(pPPO1:TP^PPO1^PPO2*) plants was indistinguishable from wild type with regard to pigmentation and development, and clearly differed from the partial complementation observed in *ppo1*(pPPO1:PPO2) individuals (Figure 5D). Interestingly, besides the complete complementation of *ppo1*, several of the *p35S:TP^PPO1^PPO2* lines containing the stronger p35S promoter displayed necrotic lesions. Immunoblot analyses revealed the affected lines to represent the strongest overexpressors of the *TP^PPO1^PPO2* fusions. Therefore, analyses presented here are based either on lines carrying pPPO1:TP^PPO1^PPO2 or weaker expressors of p35S:TP^PPO1^PPO2. Identical phenotypes were obtained when the chloroplast transit peptide of RBCS [the small subunit of ribulose bisphosphate carboxylase/oxygenase (RuBisCO)] was fused to PPO2 (TP^RBCS^PPO2).

Purification of chloroplasts and subsequent separation of membrane fractions from transgenic lines harbouring the chloroplast transit peptide fusion constructs revealed that both TP^PPO1^PPO2 and TP^RBCS^PPO2 accumulated primarily in the thylakoid fraction (Figure 5E). Immunodetection of the chloroplast envelope protein DNADJ12 (At3g15110, Pulido and Leister, 2017) and the LHCB1 signal on the Ponceau-stained membrane proved a successful separation of envelope and thylakoid membranes. The sizes of the mature fusion proteins were similar to that of the overexpressed PPO2, indicating that the fused transit peptides were the successful removed.

Quantification of TPS intermediates and end-products by HPLC supported the full complementation of the mutant phenotype by *ppo1*(p35S:TP^PPO1^PPO2) (Figure 5F). In contrast to the case in *ppo1*(p35S:PPO2), the accumulation of tetrapyrroles in homozygous *ppo1* plants expressing PPO2 fused to a cleavable chloroplast transit peptide is indistinguishable from that seen in wild-type plants (Figure 5F).

Thus, when fused to known chloroplast transit sequences, PPO2 is able to fully compensate for the loss of PPO1 even at moderate expression levels. Since even a strong overexpression of envelope-targeted, unmodified PPO2 is not able to provide a similar degree of *ppo1* complementation, the integration of Arabidopsis PPO2 into chloroplast envelope membranes can be concluded to be highly specific, supporting the assumption of a strictly separated translocation mechanism.

## Discussion

Screening of numerous publicly available T-DNA insertion lines enabled the identification of two Arabidopsis *ppo2* mutants that exhibited significant alterations in *PPO2* expression. The T-DNA insertion in *ppo2-1* is located in intron 13, and results in a reduction of *PPO2* expression. In *ppo2-2*,the integration of T-DNA was accompanied by a deletion that included part of exon 13, which effectively knocked out the gene. Consequently, no *PPO2* transcripts are detectable in the *ppo2-2* mutant (Figure 1A-F). Neither of the mutants showed any visible phenotype when grown under standard conditions, and amounts of TPS intermediates or end-products were unchanged (Figure 1 F-G). Quantification of *in-vitro* PPO activity in total leaf extracts of light-grown seedlings revealed a small, but significant decrease in enzymatic activity in the *ppo2-2* mutant in comparison to the wild type (Figure 1I). The relatively low contribution of PPO2 to overall PPO activity is compatible with the dominant expression of the PPO1 isoform under photoautotrophic conditions (Winter et al., 2007).

Knockout of *PPO1* has previously been shown to be seedling-lethal (Zhang et al., 2014), Figure 2A). However, when *ppo1* mutants were germinated in the dark, no phenotypic differences were observed among the segregating siblings of heterozygous *ppo1* mutants (Figure 2B). This demonstrates that seed development, germination and etiolated growth do not essentially depend on PPO1 enzyme activity but can be adequately sustained by PPO2 alone. To evaluate the individual contributions of the two PPO isoforms during growth in darkness and subsequent de-etiolation, steady-state levels of transcripts and proteins involved in TPS were determined (Figure 2D). In contrast to *PPO1*, as well as *HEMA1/* GLUTR1 and *FC2*, which are expressed in a characteristically light-inducible fashion, *PPO2* expression is not significantly affected by de-etiolation. The deduced stronger contribution of PPO2 to overall PPO activity under etiolating conditions was verified by measuring PPO enzyme activity in extracts of etiolated *ppo2* seedlings (Figure 2E). A dominant impact of PPO2 on total PPO activity was also observed in root tissue (Figure 2F).

The phenotype of the *ppo1ppo2* double mutant underlined the role of PPO2 in early developmental stages (Figure 2G). While seed formation and early seedling development proceed normally in *ppo1* mutants, simultaneous knock-out of both PPO isoforms is embryo-lethal, and results in the abortion of about 25% of seeds in siliques formed by *ppo1PPO1/ppo2ppo2* individuals. However, although the data in Figure 2G demonstrate a substantial contribution of PPO2 to total PPO activity in these specific developmental stages, *ppo2-2* seedlings exhibit no visible phenotype. Hence, even complete loss of PPO2 can be adequately complemented by small amounts of PPO1, e.g. under etiolating conditions.

The observed ability to mutually compensate for the loss of the other PPO isoform, at least in non-photoautotrophic tissue, raises the issue of the degree of functional overlap between PPO1 and PPO2 in Arabidopsis. Since the increased expression of PPO1 in green tissue was hypothesized to represent the main obstacle for PPO2 to compensate the loss of PPO1, PPO2 was introduced into the *ppo1* mutant background under the control of the *PPO1* promoter. In the resulting transformants, the expression of *PPO2* in leaf tissue was increased by 5-to 11-fold relative to wild type, and was therefore similar to PPO1 transcript accumulation (Figure 3A). When the T2 offspring of heterozygous (*ppo1/PPO1*) T1 transformants were screened, about 25% of them turned out to be pale green, slow-growing seedlings (Figure 3B). Since the expression of *pPPO1:PPO2* did not fully complement *ppo1*, additional lines were generated in which *PPO2* expression was driven by the strong *CaMV 35S* promoter. The resulting *ppo1*(p35S:PPO2) seedlings displayed improved complementation when compared to *ppo1(pPPO1:PPO2*), but they nevertheless failed to fully compensate for the loss of PPO1 (Figure 3C). Immunoblot analyses confirmed the overexpression of PPO2 (Figure 3D) in both transgenic lines (Figure 3D). Determination of tetrapyrrole intermediates revealed a significant accumulation of Proto, which, due to auto-oxidation, might represent either Protogen or Proto. Downstream TPS intermediates, as well as end-products, were reduced in both lines (Figure 3E). The pale-green phenotype was accompanied by a reduction of GluTR1 (Figure 3D) and ALA synthesis rates (Figure 3F) which are assumed to represent consequences of Proto(gen) accumulation. Since no other TPS proteins involved in ALA formation (GSAAT) or GluTR regulation (GBP, FLU) are affected (Figure 3D), the observed decrease of ALA synthesis can be concluded to result from a specific effect of Proto(gen) accumulation on GluTR abundance.

The inability of overexpressed PPO2 to fully complement *ppo1* excluded the possibility that insufficient amounts of PPO2 alone prevent this isoform from compensating for PPO1 deficiency. To further elucidate possible differences between the two Arabidopsis isoforms, their subcellular localization was analysed.

### Subcellular localization of PPO2

Lermontova et al. (1997) have previously described a mitochondrial localization for PPO2 in tobacco. Since the N-terminal region of Arabidopsis PPO2 closely resembles that of the *N. tabacum* isoform (Figure 5A), we tested whether the former might also be targeted to these organelles. Mitochondria were purified from wild-type and PPO2-overexpressing Arabidopsis plants (Figure 4A, B). In both cases, no significant enrichment of PPO2 in the mitochondrial fraction was observed. Hence, a mitochondrial localization of Arabidopsis PPO2 can be ruled out.

However, subsequent analyses of purified wild-type Arabidopsis chloroplasts enabled the subcellular localization of PPO1 as well of PPO2. Chloroplast membranes were further fractionated to yield preparations enriched for envelopes and thylakoids, respectively. PPO1 was detected exclusively in thylakoid membranes (Figure 4C). PPO2, in contrast, was strongly enriched in chloroplast envelope fractions. The specificity of the PPO2 localization observed in wild type was confirmed by experiments employing *ppo2-2* mutants, as well as *PPO2* overexpressor lines (Figure 4C). The observed chloroplast envelope-specific localization of PPO2 also agrees with proteomic data published by (Froehlich et al., 2003) and (Ferro et al., 2010).

PPO activity measurements revealed that conversion of Protogen in chloroplast envelope membranes depends specifically on PPO2, since the *ppo2-2* mutation effectively eliminated the enzyme activity from these fractions (Table 1). In agreement with this finding, PPO1 was localized to thylakoids with high specificity (Figure 4C). The PPO1 distribution described here contradicts data presented by (Ferro et al., 2010), who reported the presence of PPO1 in both thylakoids and envelope membranes, based on mass spectrometric analyses. Since fractionation of plastidal membranes by sucrose-gradient centrifugation enriches for envelope membranes which are largely free of contamination by thylakoids (see LHCB1 in Figure 4C), this minor difference in the description of PPO1 distribution requires further investigation. In contrast to envelope preparations, thylakoid membrane fractions obtained by density-gradient centrifugation often contain significant amounts of envelope proteins (Matringe et al., 1992) (see TIC110 in Figure 4C). Hence, assignment of chloroplast membrane proteins exclusively to the envelope is more difficult to prove conclusively and requires additional, independent evidence (see below).

The distinctive subplastidal localizations of the two Arabidopsis PPO isoforms can in principle account for the inability of overexpressed PPO2 to complement the *ppo1* mutant. However, given that the proteins share only 25% sequence identity, a functional equivalence of both isoforms is not a matter of course. To elucidate the ability of PPO2 to fully substitute PPO1 function, the N-terminal Methionine of PPO2 was replaced by the PPO1 transit peptide sequence to enable it to be targeted to thylakoids.

The N-terminus of Arabidopsis PPO2 is typical for the majority of PPO2 sequences in dicotyledonous plants. It lacks a cleavable transit peptide, as illustrated by sequence alignments revealing highly conserved amino acids starting with a serine at position 15 (Figure 5A). Cleavable transit peptides are characterized by low sequence conservation, and typically vary in length between 20 and 150 residues (Balsera et al., 2009). Indeed, the identification of mature N-termini of Arabidopsis proteins has revealed that PPO2 begins with an acetylated alanine at position 2 (https://n-terdb.i2bc.paris-saclay.fr).

Interestingly, a small group of dicotyledonous plants (*Amaranthaceae*, see spinach and *Amaranthus palmeri* in Figure 5A), as well as all monocotyledonous plants (maize and barley in Figure 5A), encode PPO2 isoforms with a distinct N-terminal extension showing characteristics of a cleavable transit peptide. Intriguingly, these N-terminally extended PPO2 sequences additionally encode a “second” methionine aligning close to the amino terminus of Arabidopsis PPO2 (boxed in Figure 5A) which could serve as an alternative translation start. We plan to examine whether both initiation codons of these extended PPO2 sequences are used and result in different subplastidal destinations.

Since several gene annotations have been published for Arabidopsis PPO (Araport 11, (Cheng et al., 2017) the possibility of a similar N-terminal extension of PPO2 must be considered. At5g14220.4 comprises a hypothetical, prolonged PPO2 reading frame with two potential start codons, respectively situated 138 bp and 144 bp upstream of the translation initiation site referred to in this study and described by the initial gene model At5g14220.1. However, we attribute no functional relevance to these upstream ATG codons for the following reasons. First, no RT-PCR amplification of the putative *PPO2* transcripts situated upstream of the annotated 58 bp of 5’UTR was possible. Secondly, RNA-seq data deposited in Araport 11 (Cheng et al., 2017) confirm that PPO2 transcripts initiate within the annotated 5’UTR of model At5g14220.1. Thirdly, the findings reported in this study are based on *PPO2* overexpression initiating at the annotated *PPO2* start codon (At5g14220.1, Figure 5A). They agree with the changes observed in plastid envelopes of *ppo2-2* knockout mutants (Figure 4 and Table 1). In addition, the exclusive accumulation of PPO2 in chloroplast envelope fractions in wild-type *A. thaliana* (Figure 5C) strongly suggests that no differently targeted PPO2 variants are present in Arabidopsis.

### Targeting of PPO2 to thylakoids results in full complementation of *ppo1*

To conclusively test the functional equivalence of PPO2 and PPO1 in Arabidopsis, the transit peptide of PPO1 was translationally fused to a PPO2 sequence starting at the alanine at position 2 (Figure 5C). Expression of TP^PPO1^PPO2 resulted in a marked enrichment of transgenic PPO2 in thylakoid membranes (Figure 5E). Complemented *ppo1*(pPPO1:TP^PPO1^PPO2) as well as *ppo1*(p35S:TP^PPO1^PPO2) exhibited a wild-type phenotype (Figure 5D) and wild-type-like amounts of TPS intermediates and end-products, thus confirming the ability of TP^PPO1^PPO2 to fully compensate for the loss of PPO1 (Figure 5F).

Interestingly, *ppo1*(pPPO1:TP^PPO1^PPO2) lines also demonstrate that even low amounts of thylakoid-targeted PPO2 are sufficient to fully complement the *ppo1* phenotype. Since a comparable degree of complementation was not even achieved using strong p35S:PPO2-based overexpressors, the targeting of *A. thaliana* PPO2 to the chloroplast envelope membrane can be deduced to be highly specific.

Although not experimentally addressed in the present study, PPO2 is assumed to be located at the inner chloroplast envelope membrane. Its substrate is provided by soluble stromal CPO1, while Proto is further processed by chelatases localized inside the chloroplast. Hence, taking into account the fact that PPO2 is an inner-envelope membrane-bound protein without a cleavable transit peptide, its targeting is assumed to occur via a non-canonical mode of integration previously described for plastidal inner-envelope membrane proteins, such as chloroplast envelope quinone oxidoreductase (ceQORH) and inner-envelope protein 32 (IEP32) (Miras et al., 2002; Nada and Soll, 2004). For ceQORH, an internal sequence motif has been reported to be responsible for envelope integration (Miras et al., 2007).

Regardless of the detailed import route for the Arabidopsis-type PPO2 sequences from dicotyledonous plants, it is worth mentioning that, despite the lack of a visible phenotype for Arabidopsis *ppo2* knockout mutants, the assignment of plant PPO2 to the inner envelope membrane of the chloroplast is of economic and biotechnological relevance. PPO2 has recently attracted attention for several reported cases of resistance against PPO-inhibiting herbicides which are caused by PPO2 point mutations (Porri et al., 2022). Such sequence variations have been discovered repeatedly in *Amaranthus* species, in which the N-terminal region of PPO2 (and thus its subcellular distribution) differs from that found in Arabidopsis (Figure 5A). The recent description of an obviously lethal phenotype caused by severe *PPO2* mutations in *Amaranthus* (Porri et al., 2022) also deviates from the Arabidopsis *ppo2-2* phenotype described in this study and hints at fundamental differences in PPO2 function between *Amaranthaceae* and dicots encoding Arabidopsis-type PPO2 sequences.

In summary, the data presented here are an attractive starting point for the re-evaluation of the role of PPO2 in TPS in higher plants. It is important to explore the individual contributions of both PPO isoforms to the pathway during plant development up to flower and seed formation, as well as in different plant tissues and organs (seeds, flowers and roots). Based on the strong impact of Arabidopsis *ppo2-2* on PPO enzyme activity in etiolated tissue and roots, further dissection of the function and subplastidal localization of the two PPO isoforms in non-photosynthetic plastids will be of particular interest.

## Acknowledgements

The excellent technical assistance of Kersten Träder is gratefully acknowledged. Sharhzad Grundler, Jana Sarrazin and Florian Salisch contributed to the study during their student projects. We are thankful to Paul Hardy for critical reading of the manuscript. Protogen for PPO activity assays was kindly provided by BASF (Germany). We acknowledge the support by the German academic exchange service (DAAD) to A.C.C.P.

## Author Contributions

B.H. and B.G. designed the research; B.H., A.C.C.P. and S.M.S. performed the experiments. B.H. and B.G. wrote the paper.

## Figure legends

**Supplementary table 1**

List of oligonucleotides used in the present study.

